# Estrogens protect bone mass by inhibiting NAD^+^ metabolism in osteoclasts

**DOI:** 10.1101/2025.07.11.664289

**Authors:** Adriana Marques-Carvalho, Gareeballah Osman Adam, Ankita Chalke, Ana Resende-Coelho, Olivia Reyes-Castro, Aaron Warren, Luis F Grilo, Claudia CS Chini, Benjamin Stronach, Farah Alturkmani, Elena Ambrogini, Eduardo N Chini, Ha-Neui Kim, Maria Almeida

**Affiliations:** Division of Endocrinology and Metabolism, Department of Internal Medicine, University of Arkansas for Medical Sciences, Little Rock, Arkansas, USA; Center for Neuroscience and Cell Biology, Center for Innovative Biomedicine and Biotechnology University of Coimbra, Portugal; Metabolism and Molecular Nutrition Laboratory, Kogod Center on Aging, Department of Anesthesiology and Perioperative Medicine, Mayo Clinic College of Medicine, Jacksonville, Florida, USA; Department of Orthopedic Surgery, University of Arkansas for Medical Sciences, Little Rock, Arkansas, USA; Center for Musculoskeletal Disease Research, University of Arkansas for Medical Sciences, Little Rock, Arkansas, USA

**Author notes:** Corresponding Author: Address all correspondence and requests for reprints to Ha-Neui Kim or Maria Almeida, University of Arkansas for Medical Sciences, Division of Endocrinology and Metabolism, 4301 W. Markham St. #587, Little Rock, 72205-7199. E- Mail (HNK) or (MA).

**Keywords:** bone resorption, mitochondria, Nampt, Sirt3

## Abstract

Estrogens protect against bone loss by reducing osteoclast number and bone resorption, primarily via direct actions on osteoclast precursors. In these cells, estrogens attenuate RANKL-induced stimulation of mitochondrial complex I, which is crucial for ATP generation through NADH oxidation. NAD^+^ promotes redox reactions and activates NAD^+^-dependent enzymes, including the mitochondrial deacetylase SIRT3. However, the contribution of NAD^+^ to the skeletal effects of estrogens remains unknown. We show that NAD^+^ levels and SIRT3 activity are upregulated by RANKL and inhibited by 17β-estradiol (E_2_) in mouse and human osteoclast precursors. Increasing NAD^+^ or the mitochondrial NAD^+^/NADH ratio reverses the inhibitory effects of E_2_ on SIRT3 activity and osteoclastogenesis *in vitro*. Deletion of *Nampt*, a key NAD salvage enzyme, reduces NAD^+^ and prevents bone loss in ovariectomized mice. Similarly, deletion of *Sirt3* in osteoclast precursors mitigates estrogen deficiency–induced bone resorption. These findings indicate that suppression of NAD^+^ levels and mitochondrial redox metabolism by estrogens contributes to their anti-resorptive effects via inhibition of SIRT3.

## Introduction

Bone is remodeled throughout life by the coordinated actions of bone-resorbing osteoclasts and bone-forming osteoblasts. Estrogens are major contributors to bone homeostasis by restraining osteoclast number and bone resorption. Loss of estrogens at menopause increases bone resorption, leading to osteoporosis and increased fracture risk. The anti-resorptive actions of estrogens are mediated, in part, through direct actions on cells of the osteoclast lineage^1–4^. Numerous *in vitro* studies with primary osteoclast cultures from humans or mice indicate that estrogens attenuate osteoclast differentiation, activity, and lifespan^2,5^. However, the molecular mechanisms responsible for these effects of estrogens on osteoclasts remain unclear.

Osteoclasts differentiate from myeloid precursors upon stimulation with macrophage colony-stimulating factor (M-CSF) and receptor activator of NF-κB ligand (RANKL), the two indispensable cytokines for bone resorption^6^. During osteoclast differentiation, RANKL promotes mitochondria oxidative phosphorylation (OXPHOS) and biogenesis to increase energy production, to satisfy the high energy-demands of bone resorption^6,7^. We have shown earlier that estrogens decrease osteoclastogenesis by inhibiting mitochondrial complex I activity, mitochondrial respiration, and ATP levels in early osteoclast progenitors^4,8^. The activity of complex I relies on electrons donated by NADH upon oxidation to NAD^+^. In addition to OXPHOS, the NAD^+^/NADH redox couple is indispensable for many other metabolic pathways^9,10^.

NAD^+^ is produced from dietary precursors such as tryptophan (TRP), nicotinic acid (NA), nicotinamide riboside (NR) and nicotinamide mononucleotide (NMN)^11^. Besides its functions in redox reactions, NAD^+^ serves as a substrate for several enzymes, including the sirtuin family of deacetylases (SIRT1-7), which degrade NAD^+^ into nicotinamide (NAM)^12–14^. NAM can be recycled into NAD^+^ via the salvage pathway. In this pathway, NAM is transformed into NMN via nicotinamide phosphoribosyltransferase (NAMPT), the rate-limiting enzyme. NMN, in turn, is converted to NAD^+^ by nicotinamide mononucleotide adenylyltransferases (NMNAT1-3).

NAD^+^ levels and metabolism vary across subcellular compartments, depending on the presence of compartment-specific NAD^+^-consuming and recycling enzymes^15,16^. For example, the presence of different sirtuins is compartment specific. SIRT3, the primary mitochondria deacetylase, maintains mitochondrial function under pathophysiologic conditions. Using global *Sirt3* knockout mice, we have shown before that SIRT3 promotes bone resorption with estrogen deficiency^17^. However, because estrogens inhibit bone resorption both via direct effects in osteoclasts and indirectly via other cell types^2^, the cellular and molecular contributions of SIRT3 to the effects of estrogens on osteoclasts remain unclear.

Recent work in macrophages has shown that NAD^+^, synthesized *de novo* from tryptophan or recycled via the salvage pathway, promotes immune functions^18–20^. However, very little is known about NAD metabolism in osteoclasts. We have shown in cell cultures that RANKL rapidly stimulates oxidative phosphorylation and estrogen attenuates this effect^8^. Because mitochondria metabolism is dependent on NAD, we examined whether RANKL promotes NAD metabolism and whether modulation of NAD contributes to the anti-osteoclastogenic effects of estrogens.

## Results

### E_2_ decreases the NAD^+^/NADH redox ratio, NAD^+^ levels, and SIRT3 activity stimulated by RANKL in osteoclast precursors

We started by evaluating NAD^+^ and NADH levels in bone marrow macrophages (BMMs) cultures using the cycling assay^21^. This method is highly specific for NAD^+^ and NADH, with no detectable cross-reactivity with other nucleotides or NAD derivatives^22^. NAD^+^ levels were also evaluated using a commercially available assay kit. The results obtained with both methods show that addition of RANKL for 24 hours increased both NAD^+^ and NADH levels (Fig. 1a and Extended Data Fig. 1). However, the NAD^+^/NADH ratio remained unchanged (Fig. 1b), suggesting an effect of RANKL on NAD metabolism beyond its stimulation of complex I activity. E_2_ had no effect on BMMs cultures in the presence of M-CSF alone but prevented the actions of RANKL. In agreement to its inhibitory effect on complex I activity, E_2_ decreased NAD^+^ levels while increasing NADH levels, resulting in a reduced NAD^+^/NADH ratio. A similar effect of RANKL and E_2_ on NAD^+^ levels was observed in mitochondrial-enriched fractions (Fig. 1c). We next determined whether the changes in NAD^+^ could affect SIRT3. The activity of SIRT3 was increased by RANKL and inhibited by E_2_ (Fig. 1d). In contrast, *Sirt3* mRNA levels were not altered by either RANKL or E_2_ (Fig. 1e). Similar to the effects in mouse cells, RANKL increased NAD^+^ levels, and SIRT3 activity in cultures of osteoclast precursors obtained from human bone marrow (Fig. 1f-g). E_2_ blocked the effects of RANKL on all these parameters.

**Fig 1.**
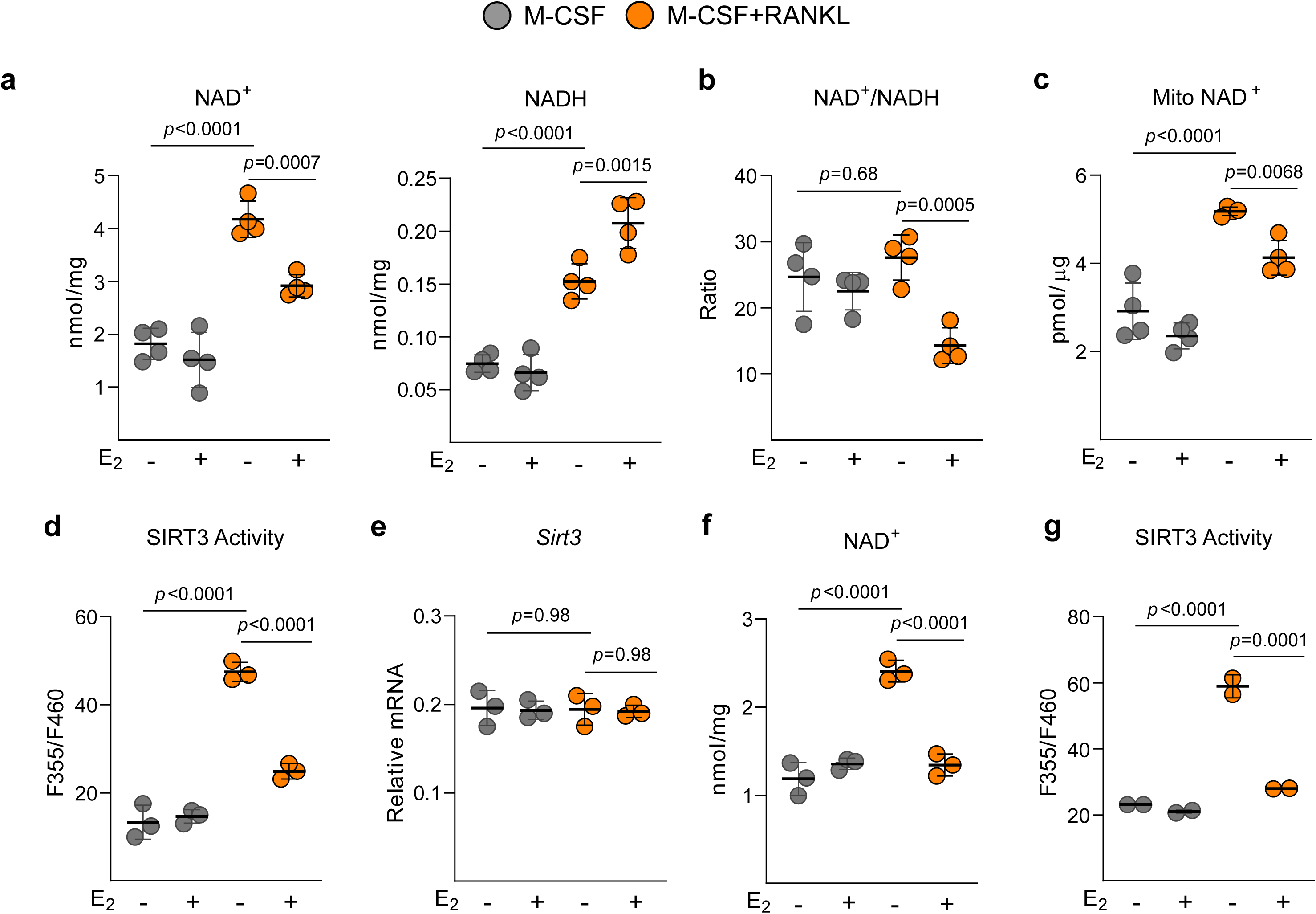
E_2_ lowers cellular NAD^+^ and disrupts the NAD^+^/NADH ratio and SIRT3 activity stimulated by RANKL. BMMs were isolated from young female C57BL/6 mice and cultured with M-CSF and RANKL in the presence or absence of E_2_ for 24 h. **(a)** NAD^+^ and NADH levels. **(b)** NAD^+^/NADH redox ratio in total or **(c)** mitochondrial-enriched fractions, **(d)** SIRT3 activity and **(e)** *Sirt3* mRNA levels. Human BMMs were isolated from femoral heads and were cultured with M-CSF and RANKL, in the presence or absence of E_2_ for 24 hours. **(f)** NAD^+^ levels and **(g)** SIRT3 activity. Line and error bars represent mean ± SD. P values determined using 2-way ANOVA followed by Šídák’s multiple comparisons test.

### Increasing NAD^+^ attenuates the anti-osteoclastogenic effects of E_2_

To examine the contribution of the decrease in NAD^+^ to the anti-osteoclastogenic effects of E_2_, we used the NAD^+^ precursor nicotinamide riboside (NR) to enhance cellular NAD^+^ levels^23^. NR contributes to NAD^+^ synthesis through direct conversion to NMN by nicotinamide riboside kinases (NRK)^24^. NR increased NAD^+^ levels and prevented the restraining effects of E_2_ on NAD^+^ levels (Fig. 2a and Extended Data Fig. 2a). NR also abrogated the inhibitory effects of E_2_ on SIRT3 activity (Fig. 2b). We next examined whether the altered NAD^+^ levels contribute to the changes caused by RANKL and E_2_ in mitochondrial respiration and ATP production. NR did not alter oxygen consumption rate and ATP levels in the absence of E_2_ but attenuated the inhibitory effect of E_2_ on both of these components of mitochondria activity (Fig. 2, c-d and Extended data Fig. 2b-f). Activation of the mitochondrial apoptosis pathway in osteoclast progenitor cells contributes to the inhibition of osteoclastogenesis by E_2_^3,4^. E_2_ increased apoptosis in osteoclast precursors, as determined by caspase 3 activity and NR also attenuated this effect (Fig. 2e). In cultures of BMMs with RANKL for 5 days, E_2_ decreased the number of TRAP positive osteoclasts (3 or more nuclei), as expected (Fig. 2f). NR modestly increased the number of osteoclasts in the absence of E_2_ and attenuated the inhibitory effects of E_2_. Consistent with the observations in mouse cells, NR prevented the inhibitory effects of E_2_ on NAD^+^ levels and the formation of osteoclasts from human BMMs (Fig. 2g-h).

**Fig 2.**
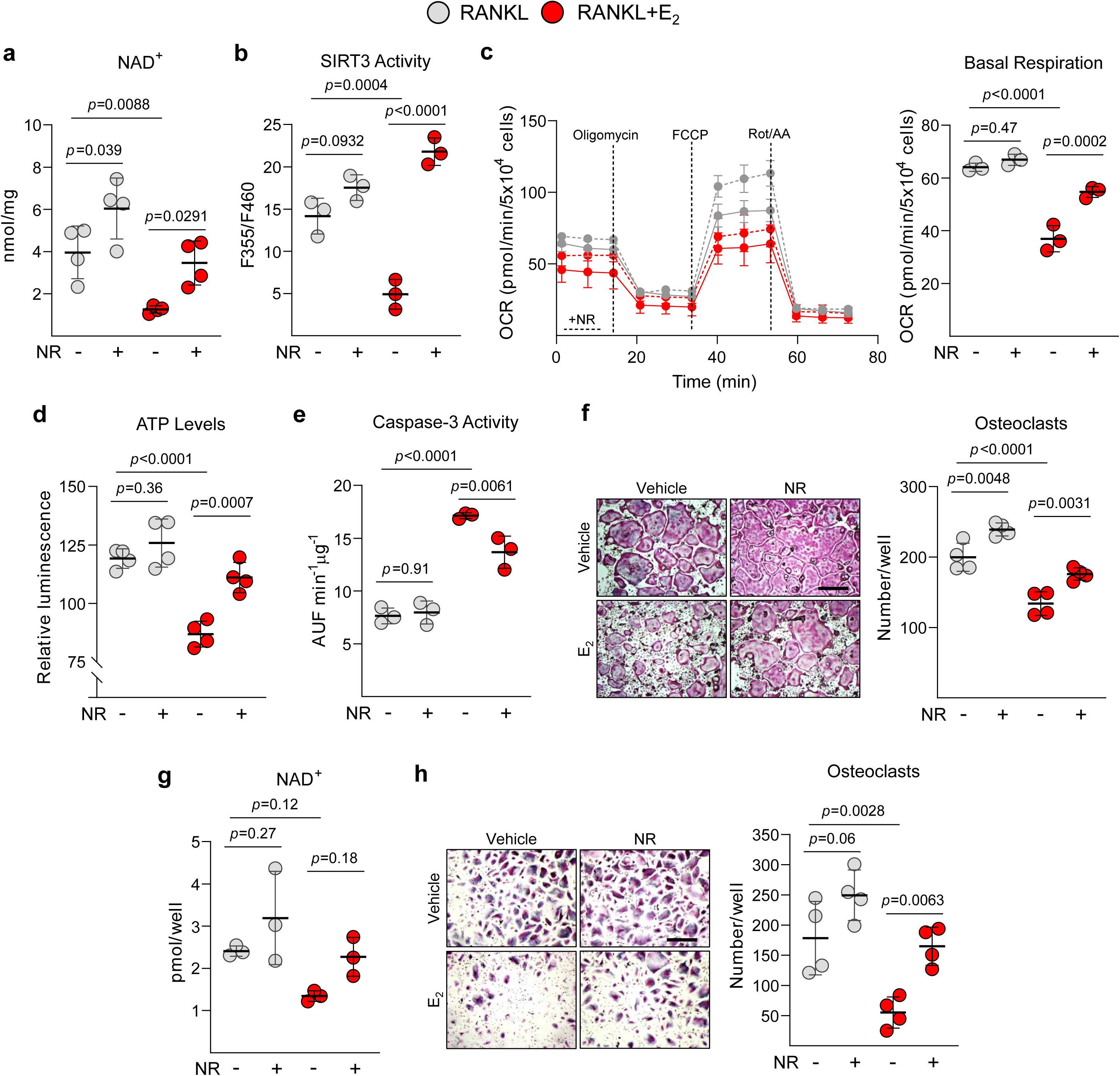
Increasing NAD^+^ restores the NAD^+^/NADH ratio and SIRT3 activity and attenuates the anti-osteoclastogenic effects of E_2_. **(a-f)** BMMs were isolated from young female C57BL/6 mice and cultured with M-CSF and RANKL in the presence or absence of E_2_ plus nicotinamide riboside (NR). **(a)** NAD^+^ levels, **(b)** SIRT3 activity, **(c)** Oxygen Consumption Rate, and **(d)** ATP levels, and **(e)** apoptosis determined by caspase-3 activity after 24 h. **(f)** Representative images and quantification of TRAP-positive osteoclasts after 5 days. **(g-h)** Human BMMs were cultured with M-CSF and RANKL in the presence or absence of E_2_ plus NR for 24 h. **(g)** NAD^+^ levels and **(h)** representative images and quantification of TRAP-positive osteoclasts after 7 days. Scale bar in F and H = 500 µm. Line and error bars represent mean ± SD. P values determined using 2-way ANOVA followed by Šídák’s multiple comparisons test.

### The NAD salvage pathway is essential for osteoclastogenesis

Given that macrophages can synthesize NAD^+^ either *de novo* from tryptophan or through the salvage pathway^20^ (Fig. 3a), we examined the contribution of these pathways to osteoclastogenesis. To this end, we utilized Phthalic Acid (PA) and FK866 as pharmacological inhibitors of Quinolinate Phosphoribosyltransferase (QPRT) and NAMPT, respectively. Treatment of BMMs with RANKL in the presence of FK866 for 24 hours decreased total NAD^+^, while PA had no effect on NAD^+^ levels (Fig. 3b and Extended Data Fig. 3a). Addition of FK866 during only the first 6 hours of differentiation was sufficient to decrease the number of TRAP-positive osteoclasts formed after 5 days, while PA had no effect on osteoclast number (Fig. 3c). FK866 also suppressed mitochondrial NAD^+^ levels and SIRT3 activity (Fig. 3d-e). In line with the lower NAD^+^, FK866 decreased mitochondrial respiration and ATP production (Fig. 3f-g and Extended Data Fig. 3b-c). In addition, FK866 increased caspase-3 activity (Fig. 3h). Interestingly, the effects of FK866 on apoptosis were greater in osteoclast precursors (4-fold) (Fig. 3h) than in myeloid precursors (<2-fold) (Extended Data Fig. 3d), suggesting that the pro-survival effects of RANKL on osteoclast precursors are highly dependent on NAD^+^. A similar inhibition on NAD^+^ levels and osteoclast formation by FK866 was observed in human-derived osteoclast cultures (Fig. 3i-j). Collectively, these results indicate that osteoclastogenesis rely on the NAD^+^ salvage pathway and that maintaining NAD^+^ levels is crucial for the stimulatory effects of RANKL on osteoclast precursors.

**Fig 3.**
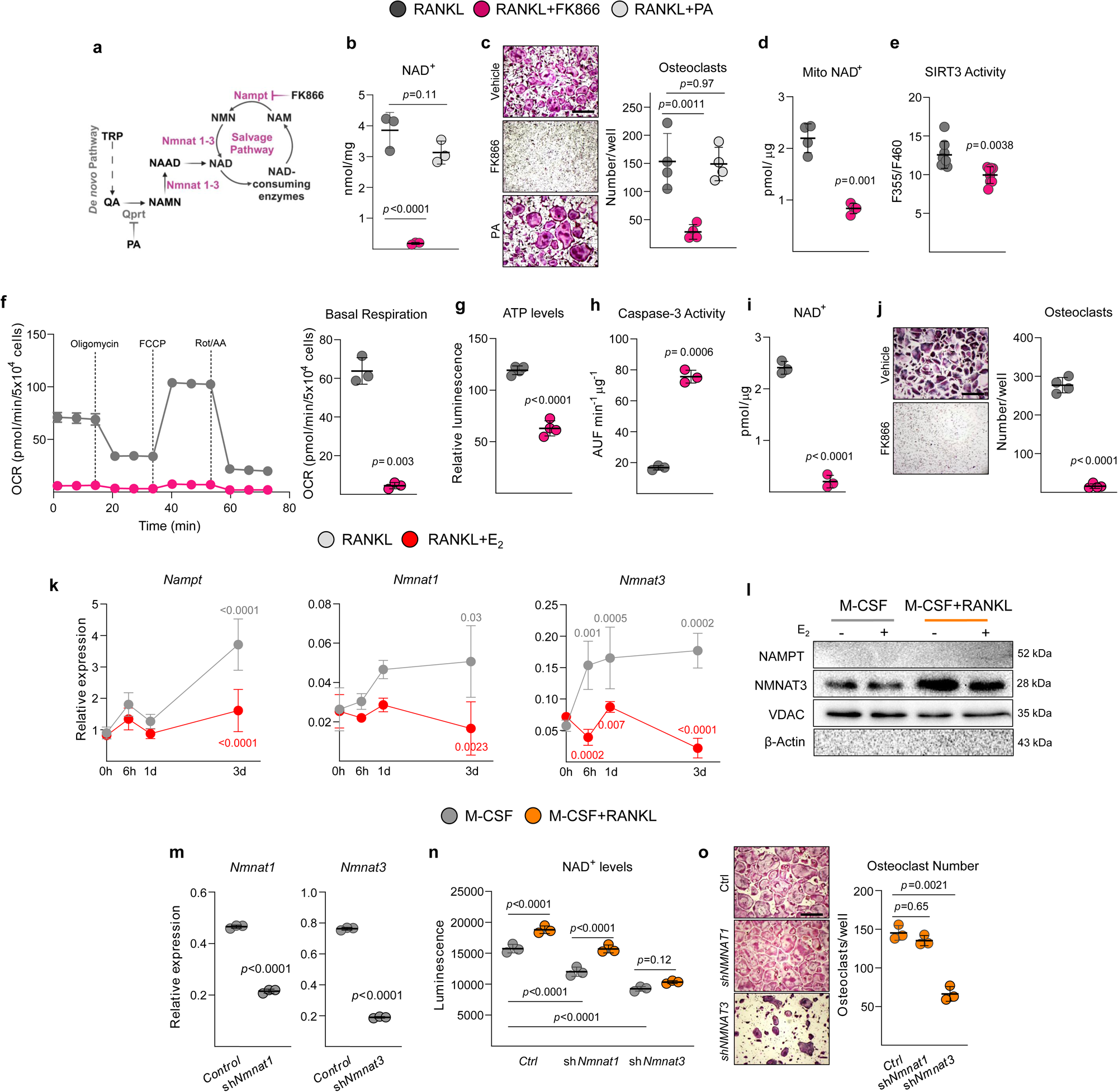
RANKL stimulates the NAD salvage pathway which is attenuated by estrogen. **(a)** Schematic illustration of NAD^+^ biosynthetic pathways. In the *de novo* pathway the essential amino acid tryptophan (TRP) from the diet is utilized to produce NAD^+^ via several reactions (dashed arrow). The salvage pathway recycles NAM, which is generated as a by-product of NAD^+^-dependent enzymes such as SIRT3; PA=Phthalic Acid. **(b-h)** BMMs were isolated from young female C57BL/6 mice and cultured with M-CSF and RANKL, in the presence or absence of FK866 or PA for 24 h. **(b)** NAD^+^ levels in whole cells. **(c)** Representative images and quantification of TRAP-positive osteoclasts after 5 days. **(d)** NAD^+^ levels in mitochondrial-enriched fractions. **(e)** SIRT3 activity. **(f)** Oxygen Consumption Rate. **(g)** ATP levels. **(h)** Apoptosis by caspase-3 activity. **(i-j)** Human BMMs were cultured with M-CSF and RANKL in the presence or absence of FK866, as above. **(i)** NAD^+^ levels after 24 h and **(j)** representative images and quantification of TRAP-positive osteoclasts after 7 days. **(k)** mRNA levels of the indicated enzymes in BMMs cultured with M-CSF and RANKL the presence or absence of E_2_ for the indicated timepoints. P values in grey indicate comparisons with time 0. P values in red indicate comparison of RANKL vs RANKL+E_2_ within the same time point. **(l)** BMMs were isolated from young female C57BL/6 mice and cultured with M-CSF and RANKL the presence or absence of E_2_ for 24 hours, representative Western blot images and quantification of the indicated proteins in mitochondrial enriched fractions; β-actin in cytosol and VDAC in mitochondria indicate efficacy of cellular compartment isolation. **(m)** mRNA levels in BMMs transduced with *Nmnat1* and *Nmnat3* sh or control sh lentivirus particles. **(n)** NAD^+^ levels after 24 h and **(o)** representative images and quantification of TRAP-positive osteoclasts after 5 days. Scale bar 500 µm. Line and error bars represent mean ± SD. P values using two-tailed Student’s t-test, 1-way ANOVA followed by Dunnett’s multiple comparisons test, or 2-way ANOVA followed by Šídák’s multiple comparisons test.

### E_2_ blunts the increased expression of NAD^+^ salvage enzymes by RANKL

In view of the requirement for the NAD^+^ salvage pathway during osteoclastogenesis, we examined whether RANKL and E_2_ altered the expression of NAMPT and NMNAT1-3. The mRNA levels of *Nmnat1* and *Nmnat3* were increased by RANKL after 24 h and *Nampt* was increased after 3 d (Fig. 3k). E_2_ inhibited the upregulation of all three enzymes. *Nmnat2* was undetectable (data not shown). NMNAT3, but not NMNAT1 or NAMPT, was found in mitochondrial-enriched fractions (Fig. 3l), in line with evidence from other cell types^25^. RANKL increased NMNAT3 expression in the mitochondria and E_2_ prevented this effect. We also searched for genes associated with NAD metabolism in our previously reported transcriptomic analysis of osteoclast differentiation ^4^. Besides the up-regulation of enzymes from both the salvage and *de novo* pathways, the gene expression of several NR and NMN transporters was upregulated upon osteoclast differentiation (Extended Data Fig. 4).

We next used short hairpin RNA (shRNA) plasmids targeting *Nmnat1* and *Nmnat3* to reduce their expression in BMMs and examine their function in osteoclasts (Fig. 3m). Knockdown of either *Nmnat1* or *Nmnat3* led to a reduction in NAD^+^ levels in BMMs compared to control shRNA-transduced cells, but only *Nmnat3* knockdown inhibited the RANKL-induced increase in NAD^+^ (Fig. 3n). In line with these findings, silencing *Nmnat3*, but not *Nmnat1*, reduced osteoclast numbers after 5 days of culture (Fig. 3o). Combined with the results obtained with FK866, these findings indicate that NAMPT and NMNAT3 are key enzymes for NAD^+^ production during osteoclastogenesis.

### Restoring mitochondrial NAD^+^/NADH redox ratio mitigates the anti-osteoclastogenic actions of E_2_

The mitochondrial electron transport chain performs two critical and interconnected processes: the oxidation of NADH to NAD^+^ by complex I and the simultaneous proton movement across the inner mitochondrial membrane to generate ATP. A balanced NAD^+^/NADH redox ratio is essential not only for ATP synthesis but also for various other metabolic processes^9,10^. To decouple the NAD^+^/NADH redox ratio from the ATP production, we employed recombinant NADH oxidase from *Lactobacillus brevis* (*Lb*NOX), which catalyzes NADH to NAD^+^ while simultaneously reducing O_2_ to H_2_O (Fig. 4a). *Lb*NOX forces an increase of the NAD^+^/NADH ratio in a compartment-specific manner, either in the cytosol (via *Lb*NOX) or the mitochondria (via mito*Lb*NOX)^26^. The efficacy of the transduction of the different plasmids was evaluated by the expression of a FLAG-tag present in the constructs. *Lb*NOX was found in the cytosolic fraction, whereas mito*Lb*NOX was found in the mitochondria fraction (Fig. 4b). The intracellular NAD^+^/NADH ratio was increased in cells expressing *Lb*NOX or mito*Lb*NOX, while ATP levels remained unaffected (Fig. 4c-d). Mito*Lb*NOX caused a higher increase in the ratio, possibly because most NADH within the cell is localized in the mitochondria. Mito*Lb*NOX but not *Lb*NOX prevented the inhibitory effects of E_2_ on the NAD^+^/NADH ratio and SIRT3 activity (Fig. 4e-f). Neither *Lb*NOX nor mito*Lb*NOX altered osteoclast differentiation (Fig. 4g). However, mito*Lb*NOX prevented the effects of E_2_ on osteoclastogenesis, while *Lb*NOX had a milder effect. These results suggest that changes in the NAD^+^/NADH ratio contribute to the anti-osteoclastogenic effect of E_2_.

**Fig 4.**
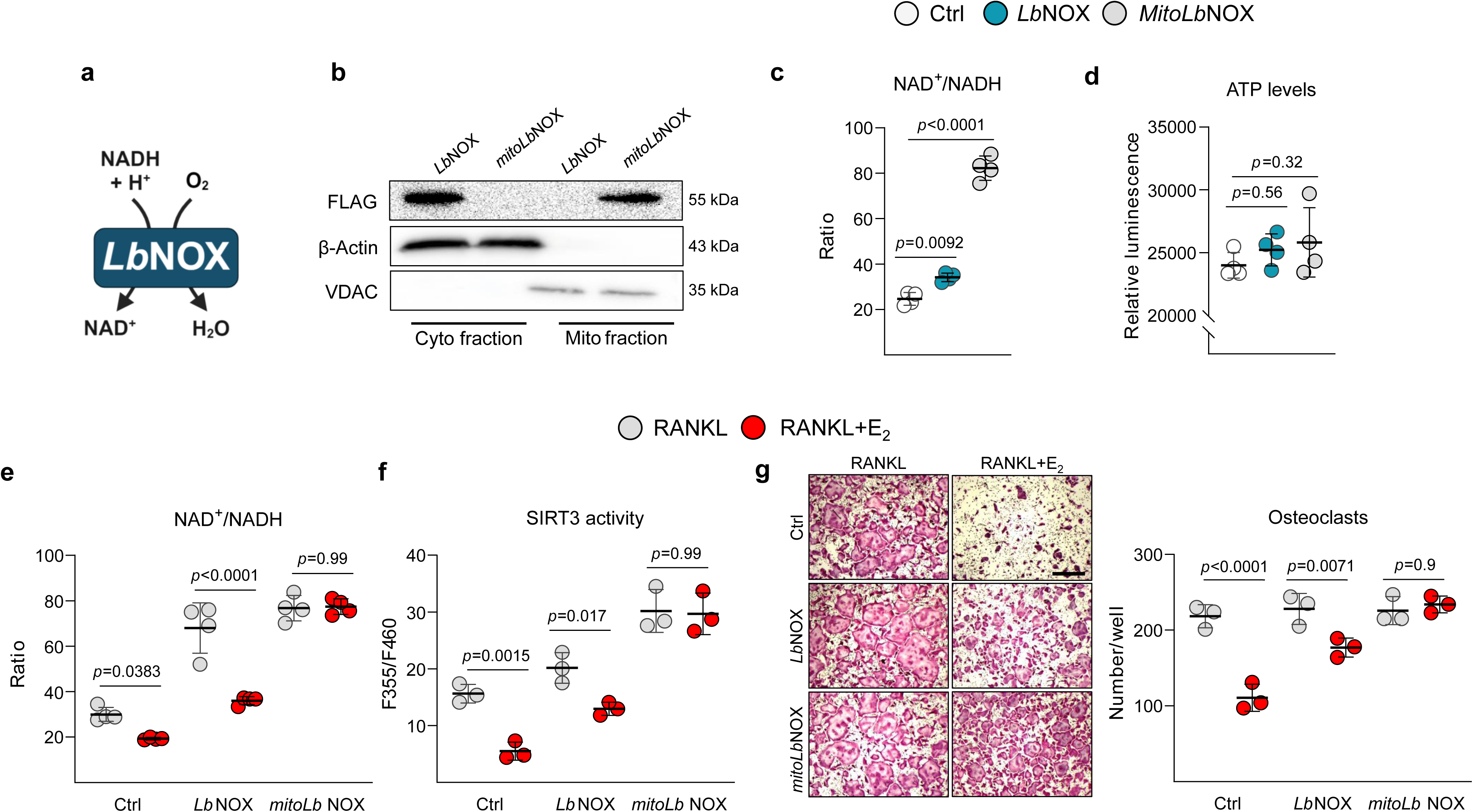
Increasing mitochondrial NAD^+^/NADH ratio attenuates the anti-osteoclastogenic effects of E_2_. **(a)** Chemical reaction catalyzed by *Lb*NOX. **(b)** Western blot for FLAG, β-actin, and VDAC from BMMs cultured with M-CSF expressing the FLAG-tagged cytosolic and mitochondrial (mito) *Lb*NOX. β-actin in cytosol and VDAC in mitochondria indicate efficacy of cellular compartment isolation. **(c)** NAD^+^/NADH redox ratio and **(d)** ATP levels in BMMs expressing *Lb*NOX or mito*Lb*NOX. **(e-g)** BMMs expressing *Lb*NOX or mito*Lb*NOX cultured with M-CSF and RANKL in the presence or absence of E_2_ (10^-8^ M) for 24 hours. **(e)** NAD^+^/NADH redox ratio. **(f)** SIRT3 activity. **(g)** Representative images and quantification of TRAP-positive osteoclasts after 5 days. Scale bar 500 µm. Line and error bars represent mean ± SD. P values by 1-way ANOVA followed by Dunnett’s multiple comparisons test or 2-way ANOVA followed by Šídák’s multiple comparisons test.

### Conditional deletion of Nampt in osteoclastic cells attenuates the loss of bone mass caused by estrogen deficiency

To elucidate the contribution of the salvage pathway to the effects of estrogens on bone resorption, we crossed mice harboring a floxed *Nampt* allele (*Nampt^f/f^*) ^27^ with mice expressing the Cre recombinase under the control of regulatory elements of the *LysM* gene, which is expressed in macrophages and neutrophils^28^ to generate *Nampt^+/f^;LysM-Cre* mice. We chose to use mice haploinsufficient for *Nampt* to avoid potential major effects on bone resorption that could compromise the interpretation of the effects of estrogen deficiency. The effectiveness of *Nampt* deletion was demonstrated by a decrease in NAMPT protein levels in cultured macrophages (Fig. 5a). *Nampt* deletion blunted the increase in NAD^+^ levels and SIRT3 activity by RANKL seen in cells from littermate controls (*Nampt^+/f^*) (Fig. 5b-c). In line with the reduced NAD^+^, respiration and ATP levels were diminished with *Nampt* deletion (Extended Data Fig. 5a-b). Similar to the findings with FK866, cells from *Nampt^+/f^;LysM-Cre* mice had increased apoptosis and formed less osteoclasts in culture (Extended Data Fig. 5c-d). These results confirm that osteoclastogenesis *in vitro* is critically dependent on the NAD salvage pathway. Nonetheless, femoral and spine BMD were unaffected in estrogen-sufficient *Nampt^+/f^;LysM-cre* mice (Fig. 5d), indicating that NAD^+^ produced by sources other than the salvage pathway compensate for the reduction of Nampt in osteoclasts *in vivo*.

**Fig 5.**
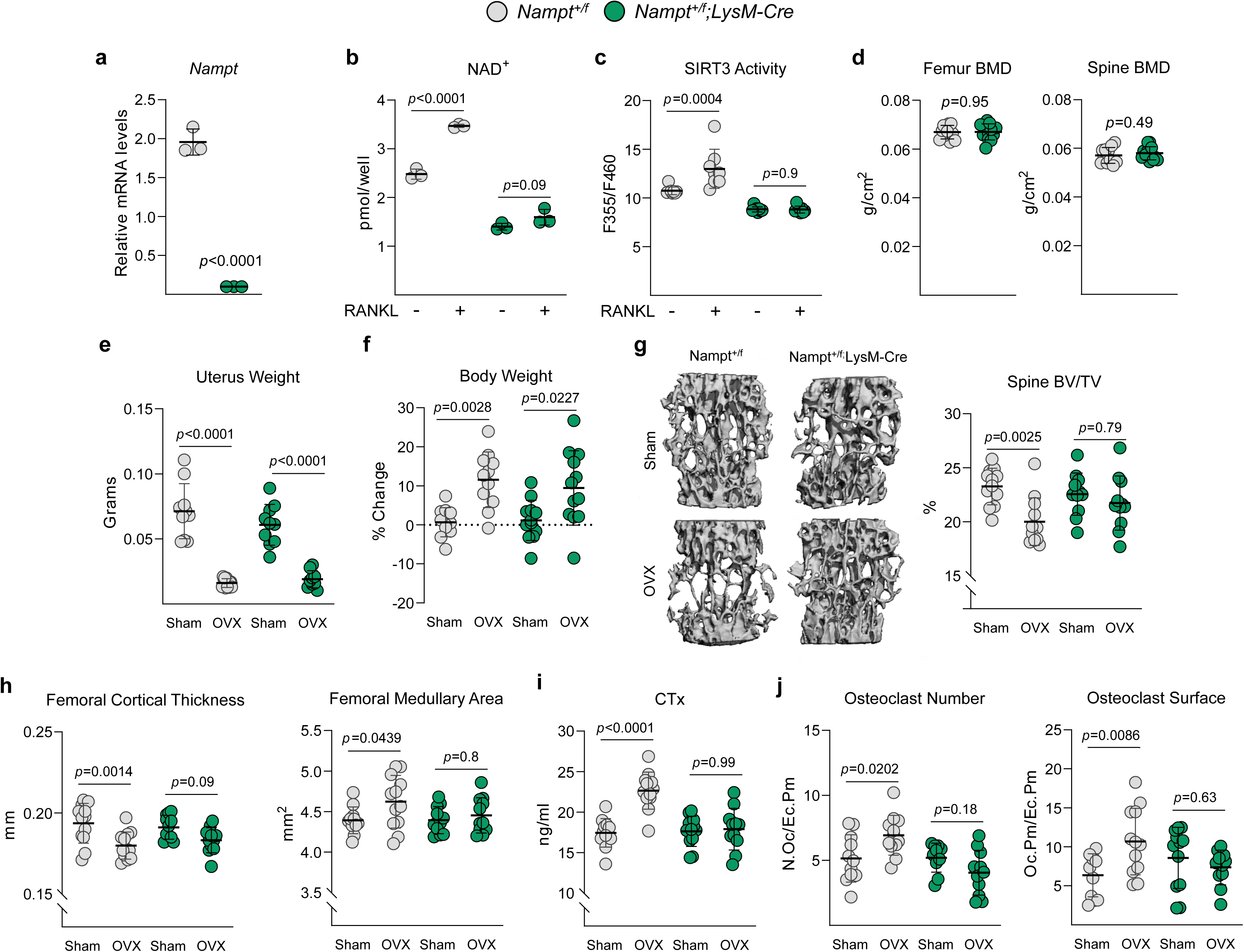
Conditional deletion of Nampt attenuates bone loss caused by estrogen deficiency. **(a)** *Nampt* mRNA levels in BMMs cultures. **(b)** NAD^+^ levels and **(c)** SIRT3 activity determined in BMMs cultured with M-CSF and RANKL for 24 hours. **(d)** BMDs by DXA in 5-month-old (n = 11–12/group) mice. **(e-j)** Five-month-old female Nampt^+/f^;LysM-Cre and Nampt^+/f^ littermate mice were sham operated or ovariectomized (OVX) and sacrificed 6 weeks later (n=11-12 animals/group). **(e)** Uterine weight at sacrifice and **(f)** percentage change of body weight determined 1 day before surgery and before sacrifice. **(g)** Micro-CT measurements of trabecular bone at the L5 vertebrae and **(h)** cortical bone at the femoral metaphysis. **(i)** Serum concentration of a collagen degradation product (CTx). **(j)** Number of osteoclast and osteoclast surface per endocortical bone perimeter at the femoral metaphysis of nondecalcified femur sections stained for TRAPase activity. Line and error bars represent mean ± SD. P values determined using two-tailed Student’s t-test and 2-way ANOVA followed by Šídák’s multiple comparisons test.

*Nampt^+/f^;LysM-Cre* and littermate controls were sham-operated or ovariectomized (OVX) at 5 months of age, and the effects of estrogen loss were assessed 6 weeks post-surgery. The efficacy of OVX was confirmed by the expected loss of uterine weight (Fig. 5e) and gain of body weight (Fig. 5f) in both control and *Nampt^+/f^;LysM-Cre* mice. Micro-CT analysis of lumbar vertebrae revealed that OVX caused loss of cancellous bone volume (Fig. 5g) due to a decrease in trabecular thickness (Extended Data Fig. 6) in control mice. However, the effect of OVX in cancellous bone of *Nampt^+/f^;LysM-Cre* mice was significantly attenuated (Fig. 5g and Extended Data Fig. 6). Likewise, OVX caused a decrease in cortical thickness at the femur in control mice due to an increase in medullary area (Fig. 5h). Deletion of *Nampt* attenuated these effects. In agreement with these findings, the serum resorption marker C-terminal telopeptide of type 1 collagen (CTx) was elevated with OVX in control but not in *Nampt^+/f^;LysM-Cre* mice (Fig. 5i). Furthermore, histomorphometric analysis of the endocortical surface of femoral bone sections revealed an increase in the number and surface covered by osteoclasts in control mice following OVX (Fig. 5j). This increase was abrogated in *Nampt^+/f^;LysM-Cre* mice.

### Deletion of Sirt3 in osteoclastic cells attenuates the bone loss caused by estrogen deficiency

Finally, we examined the functional relevance of osteoclast SIRT3 to the loss of bone mass with OVX. Mice lacking both *Sirt3* alleles under the control of *LysM-Cre* were generated and analyzed as described above. Cultured macrophages from *Sirt3^f/f^;LysM-Cre* mice had minimal if any expression of SIRT3 protein compared to cells from *LysM-Cre* littermate controls (Fig. 6a). SIRT3 activity was greatly reduced in cultured macrophages from *Sirt3^f/f^;LysM-Cre* mice (Fig. 6b). Femoral and spine BMD were not affected by *Sirt3* deletion in estrogen-sufficient mice (Fig. 6c). Following OVX, mice of both genotypes exhibited low uterine weigh and increased body weight compared to their respective sham-operated counterparts (Fig. 6d-e).

**Fig 6.**
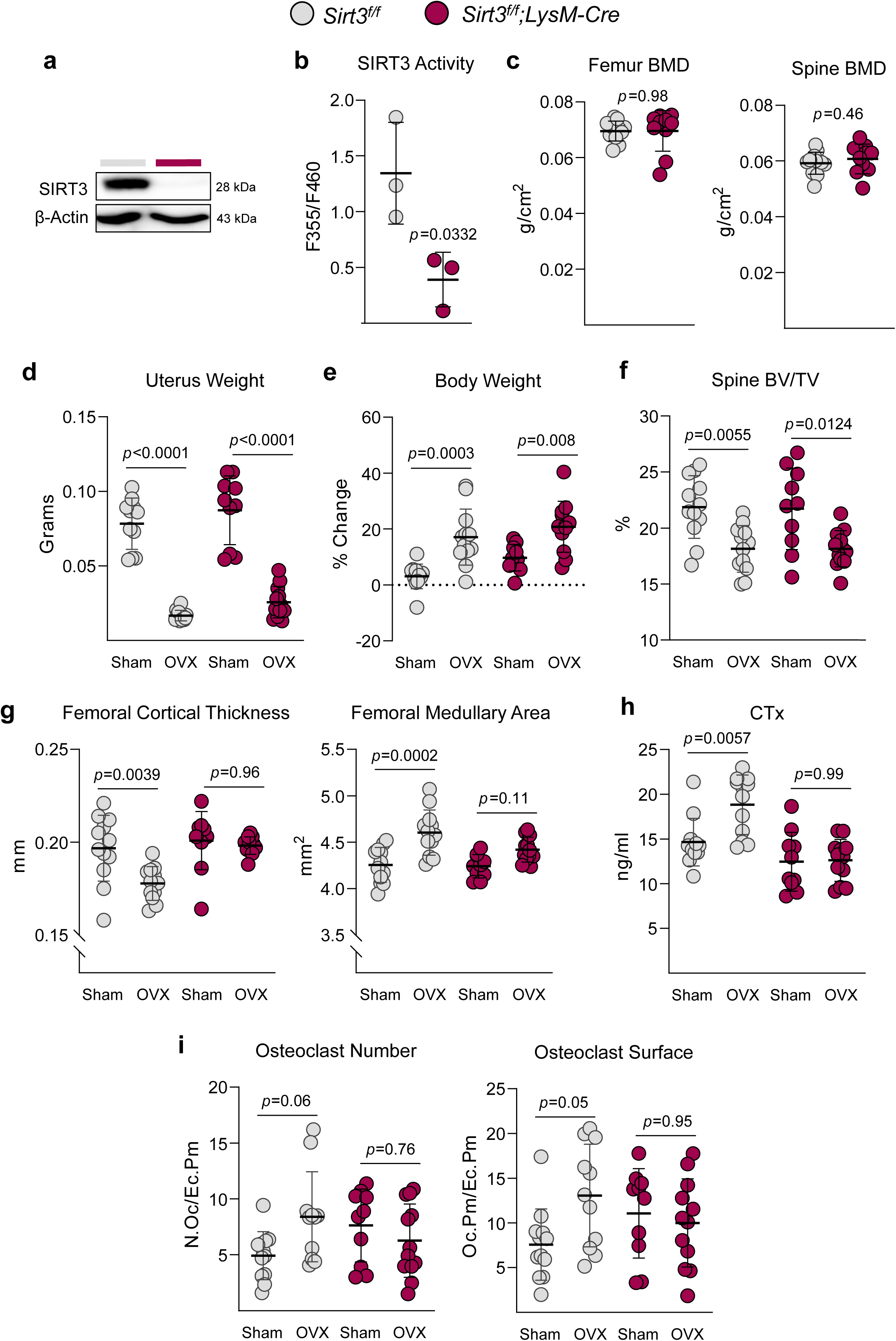
Conditional deletion of Sirt3 attenuates bone loss caused by estrogen deficiency. **(a)** Representative images of Western blots of BMMs cultures. **(b)** SIRT3 activity determined in BMMs. **(c)** BMDs by DXA in 5-month-old (n = 10–12/group) mice. **(d-i)** Five-month-old female Sirt3^f/f^;LysM-Cre and Sirt3^f/f^ littermate mice were sham operated or ovariectomized (OVX) and sacrificed 6 weeks later (n=10-12 animals/group). **(d)** Uterine weight at sacrifice and **(e)** percentage change of body weight determined 1 day before surgery and before sacrifice. **(f)** Micro-CT measurements of trabecular bone at the L5 vertebrae and **(g)** cortical bone at the femoral metaphysis. **(h)** Serum concentration of a collagen degradation product (CTx). **(i)** Number of osteoclast and osteoclast surface per endocortical bone perimeter of nondecalcified femur sections stained for TRAPase activity. Line and error bars represent mean ± SD. P values determined using two-tailed Student’s t-test and 2-way ANOVA followed by Šídák’s multiple comparisons test.

Micro-CT analysis of lumbar vertebrae showed that OVX caused a similar reduction in cancellous bone volume in both controls and *Sirt3^f/f^;LysM-Cre* mice (Fig. 6f). Additionally, OVX decreased trabecular number and thickness (Extended Data Fig. 7), and increased trabecular separation (Extended Data Fig. 7) in both groups. In contrast, the decrease in cortical thickness at the femur caused by OVX was attenuated in *Sirt3^f/f^;LysM-Cre* mice due to an decrease in medullary area (Fig. 6g). Likewise, the increase in serum CTx was attenuated by *Sirt3* deletion (Fig. 6h). Furthermore, histomorphometric analysis of the endocortical surface of femoral bone sections revealed an increase in the number and surface covered by osteoclasts in control mice following OVX (Fig. 6i). This increase was attenuated in *Sirt3^f/f^;LysM-Cre* mice.

## Discussion

Despite the evidence that cells of the osteoclast lineage are a major direct target of estrogen action^2^, until recently few molecular mechanisms have been proposed as responsible for these actions. These include suppression of RANKL-induced activator protein-1 (AP-1)-dependent transcription and Jun N-terminal kinases (JNK) and NF-kB activation^29–31^, as well as stimulation of Fas ligand (FasL) transcription and cell death^32^. Nonetheless, the relevance of these mechanisms to the bone protective actions of estrogens is lacking, and a functional role for FasL in this process could not be confirmed^4,32–38^. Using mouse genetics, as well as untargeted transcriptomic approaches, we have recently found that mitochondria is a central nexus of the actions of E_2_ in osteoclasts^4,8^. The studies presented here reveal that RANKL increases NAD^+^ levels in osteoclasts and that this is required for the increase in bone resorption upon estrogen deficiency. Furthermore, we show that attenuation of NAD^+^ metabolism contributes to the anti-osteoclastogenic effects of estrogens.

Macrophages use both the salvage and *de novo* pathway to produce NAD^+^ for immune function with inflammation^18–20^. Our findings indicate that RANKL stimulates the NAD^+^ salvage pathway and that NAD^+^ salvage is required for osteoclastogenesis *in vitro*. In contrast, the *de novo* pathway is not a major source of NAD^+^ for osteoclastogenesis. Within the skeleton, NAMPT in growth plate chondrocytes is indispensable for endochondral bone development^39^. In contrast, NAMPT in cells of the osteoblast lineage is dispensable for bone development and growth ^39^. The higher reliance of the growth plate on NAMPT is, most likely, due to the absence of vascularization and, therefore, a more limited access to NAD precursors in the circulation. In the present work, inhibition of the NAD salvage pathway using FK866 or cells from *Nampt^+/f^;LysM-Cre* mice had a dramatic effect on osteoclastogenesis *in vitro,* indicating that the NAD^+^ precursors present in the medium are rapidly exhausted during early osteoclastogenesis. However, osteoclast number and bone resorption were not impacted in estrogen-sufficient conditions in *Nampt^+/f^;LysM-Cre* mice. This is most likely due to the presence of blood vessels at sites of bone remodeling which provide a continuous supply of NAD^+^ precursors such as NR and NA. NR uptake relies on equilibrative nucleoside transporters (ENT) family members ENT1 (SLC29A1), ENT2 (SLC29A2), and ENT4 (SLC29A4). NR is then converted to NMN and transformed into NAD by NMNATs, bypassing the need for NAMPT. Further support for our premise is provided by the findings that *Slc29a1*, *Slc29a2*, and *Nmnat*1 and 3 are up-regulated during osteoclastogenesis indicating that the increase in NAD^+^ levels can occur via multiple pathways. The difference between the *in vitro* and *in vivo* results highlights the limitations of *in vitro* systems in reflecting the metabolic needs of cells *in vivo*.

The lack of a bone phenotype in *Nampt^+/f^;LysM-Cre* mice notwithstanding, our results indicate that upregulation of NAD^+^ recycling is required for the increase in bone resorption caused by estrogen deficiency. Furthermore, our results show that E_2_ prevents the increase in NAD^+^ levels, mitochondrial NAD^+^/NADH redox ratio, and SIRT3 activity caused by RANKL. Previous work of ours has elucidated an inhibitory action of E_2_ on the activity of mitochondria complex I of osteoclast precursors^4,8^. Similar to the present findings, the actions of E_2_ are seen in the presence but not in the absence of RANKL. Inhibition of complex I activity leads to a decrease in NAD^+^/NADH ratio. Thus, the changes in the redox ratio are, most likely, secondary to the changes in complex I activity.

The significance of the changes on NAD metabolism caused by RANKL and E_2_ is underscored by the fact that NAD controls hundreds of reactions throughout the cell. Interconversion of NAD^+^ and NADH fuels all paths that generate ATP, such as the mitochondrial electron transport chain, glycolysis, β-oxidation and the tricarboxylic acid (TCA) cycle. We show in the present work that inhibition of NAMPT activity decreases mitochondria respiration and ATP levels. Both fatty acid and glucose oxidation increase early during osteoclastogenesis and are needed for the production of ATP for osteoclast formation and function^40^. The rapid increase in NAD^+^ by RANKL most likely precedes and is required for these metabolic changes. The suppressive effects of E_2_ on NAD^+^ seem to be particularly relevant in mitochondria, based on the attenuation of mitochondria-localized NMNAT3 and the findings using mito*Lb*Nox. While we have not examined other metabolic pathways, it is likely that attenuation of NAD^+^ levels by estrogen in mitochondria impacts fatty acid oxidation. Inhibition of mitochondria fatty acid oxidation via deletion of carnitine palmitoyltransferase 1 or 2 (Cpt1 or Cpt2) in osteoclastic cells decreases bone resorption and increases bone mass in female, but not male, mice^41,42^. In addition, the evidence that cytoplasmic *Lb*NOX attenuated the anti-osteoclastogenic effects of estrogen suggests that redox reactions outside the mitochondria also contribute. Glucose uptake increases during osteoclast differentiation and is associated with enhanced OXPHOS but also with increased glycolysis^43^. Deletion of *Glut*1, the most abundant glucose transporter in the osteoclast lineage, impairs glycolysis, but not OXPHOS, and decreases bone resorption only in female mice^43^. The reasons for the higher dependency of the individual energy substrates in females versus males are unknown. However, the attenuation of the early up-regulation of NAD^+^ by estrogens might help explain these differences. Indeed, by restraining NAD^+^, estrogens might compromise the activity of different metabolic pathways simultaneously and, thereby, reduce metabolic flexibility in osteoclasts. Future studies should elucidate whether E_2_ inhibits osteoclastogenesis by altering several metabolic pathways secondary to the suppression of NAD^+^.

The results of our experiments using mito*Lb*Nox indicate that the decreased mitochondria NAD^+^/NADH ratio suppresses SIRT3 activity and contributes to the anti-osteoclastogenic effect of E_2_. These findings support the idea that the redox ratio per se can regulate sirtuins. It has been estimated that the levels of free NAD^+^ in the mitochondria are very similar to the Michaelis constant (*K*_m_) of SIRT3 suggesting that this enzyme is regulated by local NAD^+^ fluctuations^44^. Importantly, we show herein that SIRT3 in osteoclasts contributes to the increased bone resorption caused by estrogen deficiency. Besides SIRT3, several other NAD^+^-dependent enzymes, including SIRT1, SIRT6, and PARP1, influence osteoclastogenesis^45^. Thus, modulation of NAD^+^ levels by RANKL and E_2_ may alter the activity of these enzymes. However, SIRT1, SIRT6, and PARP1 inhibit osteoclastogenesis^46,47^ making it unlikely that these enzymes contribute to the excessive bone resorption caused by estrogen deficiency. In contrast to SIRT3, all these other enzymes reside in the nucleus, suggesting that changes in NAD^+^ metabolism driven by RANKL and E_2_ might not occur in all cellular compartments.

In conclusion, the work presented herein uncovers the mechanisms via which RANKL and estrogen actions in osteoclasts control NAD^+^ metabolism, which in turn is central to all catabolic and anabolic cellular processes. Notably, our results indicate that E_2_ inhibits NAD^+^ metabolism in osteoclasts from both mice and human, and that these effects contribute to the inhibition of bone resorption in mice. Extensive evidence has suggested that NAD^+^ availability often decreases in aged tissues and NAD^+^ supplementation alleviates age-associated diseases in mice^14,23,48,49^. This knowledge has led to several ongoing clinical trials to test the efficacy of NAD^+^ precursors in the treatment of aging-associated diseases^50^. The results obtained herein suggest that bone resorption should be monitored in patients receiving these precursors, as an increase in bone resorption is an undesirable outcome, particularly in aged populations.

## Methods

### Sex as a biological variable

Our study exclusively examined female mice because the disease modeled is only relevant in females.

### Animal experiments

Mice with conditional deletion of *Sirt3* in the myeloid lineage were generated by a two-step breeding strategy. Male homozygous *LysM-Cre* transgenic mice (The Jackson Laboratory; stock no. 004781) were crossed with *Sirt3 floxed (f/f)* mice (C57BL/6 genetic background) (The Jackson Laboratory; stock no. 031201) to generate mice heterozygous for the *Sirt3 floxed* allele with the *Cre* allele. Male *Sirt3^f/+^;ΔLysM* and female *Sirt3^f/f^* mice were intercrossed to generate *Sirt3^f/f^*and *Sirt3^f/f^;LysM-Cre* mice. Offspring were genotyped by PCR using the following primer sequences: *Sirt3*-flox primer #1 5’ CTG GCT TTG GGT TTA AGC AG 3’ and primer #2 5’ GGA GGC TGA GGC TAA AGA GC 3’.

*Nampt floxed (f/f)* mice^51^ (C57BL/6 genetic background), provided by Shinichiro Imai, Washington University School of Medicine, were crossed with *LysM-Cre* transgenic mice to obtain *Nampt^+/f^;LysM-Cre* mice. Offspring were genotyped by PCR using the following primer sequences: *Nampt*-flox primer #1 5’TTC CAG GCT ATT CTG TTC CAG 3’ and primer #2 5’ TCT GGC TCT GTG TAC TGC TGA 3’. Offspring from all genotypes were tail-clipped for DNA extraction at weaning (21 days) and then group-housed with same sex littermate. Genomic DNA extracted from tail samples was used for PCR-based genotyping following the protocols from the Jackson Laboratory. All mice used in this study were housed under standard laboratory conditions with a 12-hour dark, 12-hour light cycle, a constant temperature of 23 °C, and humidity of 48%. A standard rodent diet (Envigo, Teklad 22/5) containing 22% protein, 1.13% calcium, and 0.94% phosphorus was provided to mice ad-libitum. Investigators were blinded to study groups during animal handling and endpoint measurements. All procedures were approved by Institutional Animal Care and Use Committees of the University of Arkansas for Medical Sciences.

### DXA and micro-CT

Bone mineral density (BMD) measurements were performed by dual-energy X-ray absorptiometry (DXA) using a PIXImus densitometer (GE Lunar) on mice sedated with 2% isoflurane, and data were analyzed as previously described^52^. Scans of the entire left femur or lumbar spine were used for the measurement of BMD. Prior to micro-CT analysis, the bones were dissected, cleaned, fixed in Millonig’s phosphate buffer (Leica Biosystems), and gradually dehydrated in 100% ethanol. A micro-CT40 scanner from Scanco Medical was used to measure bone microarchitecture at medium resolution (12 μm isotropic voxel size) for quantitative determinations and integrated into 3-D voxel images (1024 × 1024-pixel matrices for each planar stack). For the latter, a Gaussian filter (sigma = 0.8, support = 1) was applied to all analyzed scans. For trabecular bone measurements, we used the vertebrae (L5), whereas cortical bone was measured at the left femoral diaphysis and metaphysis. Scan settings included X-ray tube potential (55 kVp), X-ray intensity (145 μA), and integration time (220 ms). The nomenclature used conforms to recommendations of the American Society for Bone and Mineral Research^53^. At a threshold of 200 mg/cm^3^ hydroxyapatite (HA), cortical dimensions were determined using 18–23 slices at the femoral mid diaphysis and using 50 slices between slices 300 and 350 at the distal metaphysis. Total and medullary area and circumference measurements were calculated from these slices. After defining endosteal and periosteal boundaries, an additional image processing script (“peel-iter =2”) was used to eliminate false voids caused by an imperfect wrap of the contours to the bone surface. Images were binarized with a threshold of 365 mg/cm^3^ HA, and overall porosity was determined with the “cl_image” script to obtain bone volume and void volume. To avoid the inclusion of osteocyte lacunae and canalicular space, void volumes < 31,104 μm3 (18 voxels) were excluded in the determination of porosity. All trabecular measurements were made by drawing contours every 10–20 slices and using voxel counting for BV/TV and sphere filling distance transformation indices, without presumptions about the bone shape as a rod or plate. Vertebral cortical bone thickness was determined on the ventral cortical wall using contours of cross-sectional images drawn to exclude trabecular bone, as described for femoral cortical bone.

### CTx ELISA

Blood was collected into 1.7 mL microcentrifuge tubes by retro-orbital bleeding. Blood was then kept on ice for 1 hour and centrifuged at 6,150g at 4°C for 10 minutes to separate serum from cells. Circulating CTx in serum was measured using a mouse RatLaps (CTx-I) ELISA kit (Immunodiagnostic Systems), according to the manufacturer’s directions.

### Bone histology

Freshly dissected lumbar vertebrae (L5) and right femurs were fixed overnight in 10% Millonig’s neutral-buffered formalin with 5% sucrose, followed by dehydration with ethanol and storage in 100% ethanol until embedding in methyl methacrylate (Sigma-Aldrich) and longitudinal sectioning (5 μm thickness). To measure osteoclast number and surface, sections were stained for the activity of tartrate-resistant acid phosphatase (TRAP) with the Leukocyte Acid Phosphatase (TRAP) Kit (Sigma-Aldrich) and counterstained with toluidine blue (Sigma-Aldrich). Briefly, deplasticized sections were stained with TRAP solution (a mixture of Fast Garnet GBC Base Solution, Sodium Nitrite Solution, Napthol AS-BI Phosphate Solution, Acetate Solution, and Tartrate Solution) for 1 hour and 45 min at 37 °C. Using the OsteoMeasure Analysis System (OsteoMetrics, Inc), the following primary measurements were made: endocortical perimeter (Ec.Pm), osteoclast number (N.Oc) and osteoclast surface (Oc.S/Ec.Pm). One section per sample was analyzed by a histopathologist blinded to the study groups. The terminology used in this study is that which is recommended by the Histomorphometry Nomenclature Committee of the American Society for Bone and Mineral Research^54^.

### Microarray

Cells were harvested for RNA isolation and microarray was performed as previously described ^4^. Quantile normalized gene expression was obtained from GSE111237 containing 25,697 quality-genes. To identify differentially expressed genes between myeloid precursors, pre-osteoclasts, and mature osteoclasts lacking ERα, the R packages limma and voom^55^ were used in RStudio version 4.0.5. Restricting the dataset to control samples, a model containing time and time^2^ covariables was used to determine genes with linear and quadratic behavior across time were identified and corrected for covariates and an unadjusted p-value ≤ 0.05 was considered as the significance threshold. For the computational analysis, Orange version 3.26.0 was used. All data were preprocessed for further analysis, including normalization to the interval [-1,1]. Data was clustered by similarity with hierarchical clustering on Euclidean distances and with average linkage.

### Murine cell culture

Bone marrow macrophages (BMMs) were obtained as described previously^4,56^, from 3-month-old female C57BL/6 mice. Briefly, whole bone marrow cells were flushed from the tibiae and femora, depleted of red blood cells with ACK buffer (0.01 mM EDTA, 0.011 M KHCO3, and 0.155 M NH4Cl, pH 7.3), and plated in α-MEM supplemented with 10% FBS and 1% penicillin/streptomycin with macrophage-colony stimulating factor (M-CSF; 30 ng/ml). Twenty-four hours later, non-adherent cells were then replated in Petri dishes with M-CSF (30 ng/ml) for 4 days to obtain BMMs, used as osteoclast precursors. BMMs were cultured in α-MEM complete medium with 30 ng/ml M-CSF and 30 ng/ml RANKL, in the presence or absence of 10^-8^ M 17β-estradiol (E_2_), 1 mM nicotinamide riboside (NR) and 10 µM FK866. To generate mature osteoclasts, BMMs were cultured for 4–5 days and were fixed with 10% neutral buffered formalin for 10 minutes and stained for TRAP, using the Leukocyte Acid Phosphatase Assay kit, following the manufacturer’s instructions (Sigma-Aldrich). A mature osteoclast was defined as a multinucleated (>3 nuclei) TRAP^+^ cell. Cells were plated at least in triplicate for all TRAP staining assays.

### Human cell culture

Human BMMs were isolated from the femoral heads of two female patients (aged 63 and 68) who underwent hip replacement surgery, with no known pathologies or medications affecting bone mass or structure. The bone marrow was flushed from the femoral head using an 18-gauge needle. After lysing red blood cells with ACK buffer, the remaining cells were plated in α-MEM complete medium supplemented with 30 ng/ml M-CSF on 100 mm tissue culture dishes. After 24 hours, non-adherent cells were harvested and further purified via Ficoll gradient centrifugation to isolate mononuclear hematopoietic cells. The isolated cells were then cultured on 100 mm Petri dishes for 6 days, after which all non-adherent cells (osteoclast progenitors) were collected and cultured with 30 ng/ml M-CSF and 100 ng/ml RANKL to induce osteoclast differentiation. A mature osteoclast was defined as a multinucleated cell with more than three nuclei and positive for TRAP staining. All TRAP staining assays were performed in triplicate.

### NAD^+^/NADH assay

Intracellular NAD^+^ and NADH levels were quantified using an enzymatic cycling assay (PMID: 27304511; PMID: 29719225: PMID: 30101155). For each condition, either 1.5x10^6^ cells (for NAD^+^) or 3x10^6^ cells (for NADH) were collected. To extract NAD^+^, cells were lysed with 100 μl of 10% trichloroacetic acid (TCA), sonicated, and centrifuged at 11,500 g for 2 min at 4°C, The supernatants containing NAD^+^ were collected, and the pellets were resuspended in 100 μl of 0.2 N NaOH for protein quantification using the BCA assay. TCA was removed by phase separation using a 2:1 ratio of organic solvent (3 parts ethyl ether to 1 part trioctylamine). The aqueous phase was collected, and the pH was neutralized with 1M Tris-HCl (pH 8). For NADH extraction, cells were lysed with 100 μl of 500 mM NaOH and 5 mM EDTA, followed by sonication and incubation at 60°C for 30 minutes. After cooling, the pH was adjusted with 5M HCl. For the cycling assay, samples were diluted in 100 mM sodium phosphate buffer (pH 8.0) and added to white 96-well plates at 100 μl per well. An equal volume of the reaction mix was added, containing 0.76% EtOH, 4 μM FMN, 27.2 U/mL alcohol dehydrogenase, 1.8 U/mL diaphorase, and 8 μM resazurin. Fluorescence (λex/em =544/590 nm) was measured over 60 minutes in a Cytation™ 5 microplate reader.

NAD^+^ levels were also measured by using the EnzyFluoTM NAD^+^/NADH Assay kit (Bioassay Systems). Briefly, BMMs were plated in a 96-well black-wall tissue culture plate and culture for 24 hours with the indicated treatments. NAD extraction buffers were then added to the respective wells according to the manufacturer instructions. The fluorescent signal (λex/em =530/585 nm) was quantified in a Cytation™ 5 microplate reader.

### Mitochondrial isolation

BMMs were cultured in 100 mm dishes for 24 hours with the indicated treatments and mitochondrial-enriched fractions were obtained using the Mitochondria isolation kit (Thermo Fisher Scientific) according to the manufacturer instructions, using 2x10^7^ cells. The pellets were resuspended to a final protein concentration of 2 mg/mL.

### Sirt3 activity

BMMs were cultured in 100 mm dishes for 24 hours with the indicated treatments and mitochondrial-enriched fractions were obtained as described above. Sirt3 activity was determined in the mitochondrial fractions using the Sirt3 Activity Assay kit (Abcam), according to the manufacturer instructions.

### Mitochondrial respiration and cellular bioenergetics

Bone marrow macrophages were plated in Seahorse XF96 plates and treated for 24 hours with the indicated treatments. The medium in the wells was replaced with XF assay medium (Agilent), and the plates were kept in a non-CO 2 incubator for 60 min at 37 °C. After recording 3 total cellular respiration measurements with the XF96 analyzer, consecutive injections of 2.5 µM oligomycin, 2 µM FCCP and 0.7 µM antimycin A/rotenone cocktail were used to determine mitochondrial basal respiration, maximal respiration ATP-linked respiration, spare respiratory capacity, proton leak and non-mitochondrial oxygen consumption.

### ATP production

Intracellular ATP levels were measured by a luciferin-luciferase based assay using CellTiter-Glo® Luminescent Cell Viability Assay (G7570, Promega), according to the manufacturer protocol. Briefly, BMMs were cultured in a 96-well white-wall tissue culture plate and treated for 24 hours with the indicated treatments. Cell culture media was replaced with 100 µl of assay reagent (CellTiter-Glo Buffer and CellTiter-Glo Substrate). Each extracted samples were mixed for 2 minutes on an orbital shaker to promote cell lysis, followed by a 10 min incubation at room temperature. The luminescence signal was monitored in a Cytation™ 5 microplate reader.

### Caspase 3 activity

Caspase-3 activity was determined by measuring the degradation of the fluorescent substrate DEVD-AFC (Biomol Research Labs, Plymouth, PA) and protein concentration was measured by a Bio-Rad detergent–compatible kit (Bio-Rad, Hercules, CA), as described previously [10]. Briefly, BMMs were cultured in 6-well plates for 24 hours with the indicated treatments and then lysed with 20 mM Tris-HCl (pH 7.5), 150 mM NaCl, 1 mM EDTA, 10 mM NaF, 1 mM sodium orthovanadate, 5 mg ml−1 leupeptin, 0.14 U ml−1 aprotinin, 1 mM phenylmethylsulfonylfluoride, and 1% Triton X-100. Cell lysates were then transferred to a new plate and incubated with 50 mM DEVD-AFC in 50 mM HEPES (pH 7.4), 100 mM NaCl, 0.1% CHAPS, 10 mM DTT, 1 mM EDTA, and 10% glycerol. The released fluorescent signal (λex/em =400/510 nm) was measured kinetically in a Cytation™ 5 microplate reader.

### Western blot analysis

BMMs were cultured in 6-well plates for 24 hours with the indicated treatments. Cells were then washed twice with ice-cold PBS and lysed with a buffer containing 20 mM Tris-HCL, 150 mM NaCl, 1% Triton X-100, and protease inhibitor mixture, and phosphatase inhibitor cocktail (Sigma-Aldrich) on ice for 30 minutes. The cell lysates were centrifuged at 13,200 rpm for 15 min at 4°C, and the supernatants were collected in a new 1.5 ml microcentrifuge tube. The protein concentration of cell lysates was determined using a DC Protein Assay kit (Bio-Rad). The extracted protein (30–40 μg per sample) was subjected to 10%–12% SDS-PAGE gels and transferred electrophoretically onto polyvinyl difluoride membranes (MilliporeSigma). The membranes were blocked in 5% fat-free milk/Tris-buffered saline for 90 min and incubated with each primary antibody overnight, followed by secondary antibodies conjugated with horseradish peroxidase. The following primary monoclonal antibodies were used: NAMPT (Abcam, ab236874, 1:1000); NMNAT3 (Santa Cruz, sc-390433, 1:100); FLAG (Cell Signaling; 2368; 1:1000); VDAC (Cell Signaling; D73D12; 1:1000) and β-actin (Santa Cruz; sc-47778; 1:5000). Bound antibodies were detected with ECL reagents (MilliporeSigma) and were imaged and quantified with a VersaDoc imaging system (Bio-Rad).

### Quantitative RT-PCR

Total RNA was purified from cultured BMMs using TRIzol reagent (Invitrogen, Carlsbad, CA). After extraction, RNA was quantified using a Nanodrop instrument (Thermo Fisher Scientific), and 2 μg of RNA was then used to synthesize cDNA using a High-Capacity cDNA Archive Kit (Applied Biosystems, Grand Island, NY) according to the manufacturer’s instructions. Transcript abundance in the cDNA was measured by qPCR using Taqman Universal PCR Master Mix (Thermo Fisher Scientific). The primers and probes for murine *Nampt* (Mm 00451938_m1); *Nmnat1* (Mm 01257929_m1); *Nmnat3* (Mm 00513791_m1) were manufactured by the TaqMan® Gene Expression Assays service (Applied Biosystems). Relative mRNA expression levels were normalized to the house-keeping gene ribosomal protein S2 (Mm00475528_m1) using the ΔCt method.

### Retrovirus Production and BMMs infection

*LbNOX* and mito*LbNOX* retroviral constructs were produced by subcloning the respective genes from pUC57-LbNOX and pUC57-mito*Lb*NOX plasmids into a pMXs-Ires-puro vector. The plasmids, pUC57-*Lb*NOX (Addgene plasmid #75285; http://n2t.net/addgene:75285; RRID:Addgene_75285) and pUC57-mito*Lb*NOX (Addgene plasmid #74448; http://n2t.net/addgene:74448; RRID:Addgene_74448) were a gift from Vamsi Mootha. Viral particle production was conducted in the Platinum-E (Plat-E) cell line using the Lipofectamine 3000 transfection reagent (Thermo Fisher), following the manufacturer protocol. Bone marrow cells were isolated from 3-month-old female C57BL/6 mice as previously described. After 24 hours, non-adherent cells were subjected to a Ficoll-Hypaque gradient, and the cells at the interface were cultured in 30 ng/ml M-CSF, 8 µg/ml polybrene, and the Plat-E supernatant containing the retrovirus vector for 16 hours. Infected BMMs were then cultured with M-CSF for an additional 24 hours, followed by puromycin selection (2 µg/ml) for 48 hours to eliminate uninfected cells.

### Lentiviral transduction of BMMs

Lentiviral particles, pLKO.1 with short hairpin RNAs (shRNAs), containing *Nmnat1* and *Nmnat3*-specific sequences (GenBank accession No. NC_000070.7 and NC_000075.7, respectively) were purchased from Santa Cruz. Non-targeted shRNA lentiviral particles (pLKO.1-empty vector) were used as shRNA control. Whole bone marrow cells were obtained from 3-month-old female C57BL/6 mice, as described above. Twenty-four hours later, non-adherent cells were submitted to a Ficoll-Hypaque gradient, and cells at the interface were cultured in the presence of M-CSF (30 ng/ml), polybrene (8 µg/ml; Santa Cruz Biotechnology), and the lentiviral particles for 16 h. We further cultured the infected BMMs with M-CSF for 24 h and then added puromycin (2 µg/ml; Santa Cruz Biotechnology) for 48 h to remove uninfected cells.

### Statistics

All *in vitro* experiments were repeated at least twice. Depending on the assay, within each experiment we used 3-10 wells per treatment. The representative experiments are shown in the figures and depict individual data point, average, and SD. Statistical analysis and graphical design were performed in GraphPad Prism 9 software (Graphpad Software). Group mean values were compared, as appropriate, by one-way ANOVA followed by Dunnett’s multiple comparisons test or two-way ANOVA followed by Šídák’s multiple comparisons test, after determining that the data were normally distributed and exhibited equivalent variances by D’Agostino & Pearson test and Shapiro-Wilk test. For experiments involving a comparison of only 2 groups, a 2-tailed Student’s t-test was used. The interaction terms of the two-way ANOVA for the *in vivo* studies can be found in Supplemental Table 1.

### Study Approvals

All animal studies were approved by the UAMS Institutional Animal Care and Use Committee (AUP 4188). Collection and de-identification of human bone samples were coordinated by the Histology, Biomechanics and Human Tissue Core of the UAMS Center for Musculoskeletal Disease Research, the UAMS Winthrop P. Rockefeller Cancer Institute Tissue Bank and Procurement Service and approved by the UAMS IRB (protocol #262940). All participants provided written, informed consent before study procedures occurred, with continuous consent ensured throughout participation.

## Data availability

Data will be made available from the corresponding author upon reasonable request. Source data are provided with this paper.

## Acknowledgements

This work was supported by the National Institutes of Health (R01AR082418, R01AR080736, R56AR056679), Center for Musculoskeletal Disease Research COBRE (P20GM125503), and the UAMS Bone and Joint Initiative. We thank Stavros Manolagas, Charles O’Brien, and Ryan Porter for critically reviewing the manuscript.

## Author contributions

AMC, HNK, and MA designed the experiments. AC and AW performed animal experimentation. GOA, ORC, and HNK performed DXA and micro-CT measurements. AMC, AC, and ARC performed histological analysis. AMC and GOA performed *in vitro* experiments. LFG analyzed microarray data. ENC and CCSC provided methodology, technical support, and data interpretation on NAD assays. EA, BS, and FA obtained human femoral heads from patients. AMC and MA wrote the manuscript. All authors reviewed the manuscript.

## EXTENDED DATA FIGURE LEGENDS

**Extended Data Fig. 1 Validation of RANKL and E_2_-Induced alterations in NAD^+^ using a commercially available assay kit.** BMMs were isolated from young female C57BL/6 mice and cultured with M-CSF and RANKL in the presence or absence of E_2_ for 24 h.

**Extended Data Fig. 2 Increasing NAD^+^ restores mitochondrial respiration.** BMMs were isolated from young female C57BL/6 mice and were cultured with M-CSF and RANKL the presence or absence of E_2_ plus nicotinamide riboside (NR) for 24 hours. **(a)** NAD^+^ levels measured using a commercially available assay kit. **(b)** ATP-linked, **(c)** Maximal, **(d)** Spare, **(e)** Proton Leak and **(f)** Non-Mitochondrial Respiration were determined by the measurement of the Oxygen Consumption Rate using the Seahorse Mitostress assay. Line and error bars represent mean ± SD. P values determined using 2-way ANOVA followed by Šídák’s multiple comparisons test.

**Extended Data Fig. 3 Reduced NAD^+^ levels in macrophages show minimal impact compared to osteoclast precursors. (a-c)** BMMs were isolated from young female C57BL/6 mice and were cultured with M-CSF in the presence or absence of FK866 for 24 hours. **(a)** NAD^+^ levels measured using a commercially available assay kit. **(b)** Oxygen consumption, **(c)** ATP levels and **(d)** Apoptosis determined by the measurement of caspase-3 activity after 24 hours. Line and error bars represent SD. P values determined using two-tailed Student’s t-test.

**Extended Data Fig. 4 NAD metabolism-associated genes in osteoclastic cells lacking ERα.** Gene expression heat map of NAD metabolism-associated genes that have an adjusted- *p* value < 0.05, in murine BMMs cultured with M-CSF for 5 days (myeloid precursors), and with M-CSF plus RANKL for 2 (pre-osteoclasts) or 4 days (mature osteoclasts). Each column represents one individual well.

**Extended Data Fig. 5 Deleting Nampt mimics E_2_ effects in vitro. (a)** Basal respiration and **(b)** ATP levels of BMMs isolated from female Nampt^+/f^;LysM-Cre and littermates Nampt^+/f^ mice and cultured with M-CSF and RANKL for 24 hours. **(c)** Representative images and quantification of TRAP-positive osteoclasts after 5 days. Scale bar 500 µm. **(d)** Apoptosis determined by the measurement of caspase-3 activity. Line and error bars represent mean ± SD. P values determined using two-tailed Student’s t-test and 2-way ANOVA followed by Šídák’s multiple comparisons test.

**Extended Data Fig 6 Effect of Nampt deletion in the microarchitecture of trabecular bone.** Five-month-old female Nampt^+/f^;LysM-Cre and littermates Nampt^+/f^ mice were sham operated or ovariectomized (OVX) and sacrificed 6 weeks later (n=10-12 animals/group). Quantification of microarchitecture of trabecular bone in L5 vertebrae. Line and error bars represent mean ± SD. P values determined using 2-way ANOVA followed by Šídák’s multiple comparisons test.

**Extended Data Fig. 7 Effect of Sirt3 deletion in the microarchitecture of trabecular bone.** Five-month-old female Sirt3^f/f^;LysM-Cre and littermates Sirt3^f/f^ mice were sham operated or ovariectomized (OVX) and sacrificed 6 weeks later (n=11-12 animals/group). Quantification of microarchitecture of trabecular bone in L5 vertebrae. Line and error bars represent mean ± SD. P values determined using 2-way ANOVA followed by Šídák’s multiple comparisons test.

## EXTENDED DATA TABLE LEGENDS

**Extended Data Table 1 Interaction p values.** Interaction p values determined using 2-way ANOVA followed by Šídák’s multiple comparisons test.

## References

1 Manolagas, S. C. Steroids and osteoporosis: the quest for mechanisms. J Clin Invest 123, 1919-1921 (2013). 68062 [pii];10.1172/JCI68062 [doi]

2 Almeida, M. et al. Estrogens and Androgens in Skeletal Physiology and Pathophysiology. Physiol Rev 97, 135–187 (2017). 97/1/135 [pii];10.1152/physrev.00033.2015 [doi]

3 Hughes, D. E. et al. Estrogen promotes apoptosis of murine osteoclasts mediated by TGF-β. Nat. Med 2, 1132–1136 (1996).

4 Kim, H. N. et al. Estrogens decrease osteoclast number by attenuating mitochondria oxidative phosphorylation and ATP production in early osteoclast precursors. Sci Rep 10, 11933 (2020). 10.1038/s41598-020-68890-7

5 Khosla, S., Oursler, M. J. & Monroez, D. G. Estrogen and the skeleton. Trends Endocrinol Metab 23, 576–581 (2012). S1043-2760(12)00052-5 [pii];10.1016/j.tem.2012.03.008 [doi]

6 Teitelbaum, S. L. & Ross, F. P. Genetic regulation of osteoclast development and function. Nat Rev Genet 4, 638–649 (2003). 10.1038/nrg1122 [doi];nrg1122 [pii]

7. Boyle, W. J., Simonet, W. S. & Lacey, D. L. Osteoclast differentiation and activation. Nature 423, 337-342 (2003). 10.1038/nature01658

8 Marques-Carvalho, A., Sardao, V. A., Kim, H. N. & Almeida, M. ECSIT is essential for RANKL-induced stimulation of mitochondria in osteoclasts and a target for the anti-osteoclastogenic effects of estrogens. Front Endocrinol (Lausanne) 14, 1110369 (2023). 10.3389/fendo.2023.1110369

9 Santidrian, A. F. et al. Mitochondrial complex I activity and NAD+/NADH balance regulate breast cancer progression. J Clin Invest 123, 1068–1081 (2013). 10.1172/JCI64264

10 Srivastava, S. Emerging therapeutic roles for NAD(+) metabolism in mitochondrial and age-related disorders. Clin Transl Med 5, 25 (2016). 10.1186/s40169-016-0104-7

11 Xie, N. et al. NAD(+) metabolism: pathophysiologic mechanisms and therapeutic potential. Signal Transduct Target Ther 5, 227 (2020). 10.1038/s41392-020-00311-7

12 Lee, C. F. et al. Normalization of NAD+ Redox Balance as a Therapy for Heart Failure. Circulation 134, 883–894 (2016). 10.1161/CIRCULATIONAHA.116.022495

13 Wei, C. C. et al. NAD replenishment with nicotinamide mononucleotide protects blood-brain barrier integrity and attenuates delayed tissue plasminogen activator-induced haemorrhagic transformation after cerebral ischaemia. Br J Pharmacol 174, 3823–3836 (2017). 10.1111/bph.13979

14 de Picciotto, N. E. et al. Nicotinamide mononucleotide supplementation reverses vascular dysfunction and oxidative stress with aging in mice. Aging Cell 15, 522–530 (2016). 10.1111/acel.12461

15 Yang, H. et al. Nutrient-sensitive mitochondrial NAD+ levels dictate cell survival. Cell 130, 1095-1107 (2007). 10.1016/j.cell.2007.07.035

16 Berger, F., Lau, C., Dahlmann, M. & Ziegler, M. Subcellular compartmentation and differential catalytic properties of the three human nicotinamide mononucleotide adenylyltransferase isoforms. J Biol Chem 280, 36334–36341 (2005). 10.1074/jbc.M508660200

17 Ling, W. et al. Mitochondrial Sirt3 contributes to the bone loss caused by aging or estrogen deficiency. JCI Insight 6 (2021). 10.1172/jci.insight.146728

18 Groth, B., Venkatakrishnan, P. & Lin, S. J. NAD(+) Metabolism, Metabolic Stress, and Infection. Front Mol Biosci 8, 686412 (2021). 10.3389/fmolb.2021.686412

19 Okabe, K., Yaku, K., Tobe, K. & Nakagawa, T. Implications of altered NAD metabolism in metabolic disorders. J Biomed Sci 26, 34 (2019). 10.1186/s12929-019-0527-8

20 Minhas, P. S. et al. Macrophage de novo NAD(+) synthesis specifies immune function in aging and inflammation. Nat Immunol 20, 50–63 (2019). 10.1038/s41590-018-0255-3

21 Kanamori, K. S. et al. Two Different Methods of Quantification of Oxidized Nicotinamide Adenine Dinucleotide (NAD(+)) and Reduced Nicotinamide Adenine Dinucleotide (NADH) Intracellular Levels: Enzymatic Coupled Cycling Assay and Ultra-performance Liquid Chromatography (UPLC)-Mass Spectrometry. Bio Protoc 8 (2018). 10.21769/BioProtoc.2937

22 Camacho-Pereira, J. et al. CD38 Dictates Age-Related NAD Decline and Mitochondrial Dysfunction through an SIRT3-Dependent Mechanism. Cell Metab 23, 1127–1139 (2016). 10.1016/j.cmet.2016.05.006

23 Kim, H. N. et al. A decrease in NAD(+) contributes to the loss of osteoprogenitors and bone mass with aging. NPJ Aging Mech Dis 7, 8 (2021). 10.1038/s41514-021-00058-7

24 Bieganowski, P. & Brenner, C. Discoveries of nicotinamide riboside as a nutrient and conserved NRK genes establish a Preiss-Handler independent route to NAD+ in fungi and humans. Cell 117, 495–502 (2004). 10.1016/s0092-8674(04)00416-7

25 Covarrubias, A. J., Perrone, R., Grozio, A. & Verdin, E. NAD(+) metabolism and its roles in cellular processes during ageing. Nat Rev Mol Cell Biol 22, 119–141 (2021). 10.1038/s41580-020-00313-x

26 Titov, D. V. et al. Complementation of mitochondrial electron transport chain by manipulation of the NAD+/NADH ratio. Science 352, 231–235 (2016). 10.1126/science.aad4017

27. Lin, J. B. et al. NAMPT-Mediated NAD(+) Biosynthesis Is Essential for Vision In Mice. Cell Rep 17, 69-85 (2016). 10.1016/j.celrep.2016.08.073

28 Clausen, B. E., Burkhardt, C., Reith, W., Renkawitz, R. & Forster, I. Conditional gene targeting in macrophages and granulocytes using LysMcre mice. Transgenic Res 8, 265–277 (1999). 10.1023/a:1008942828960

29 Shevde, N. K., Bendixen, A. C., Dienger, K. M. & Pike, J. W. Estrogens suppress RANK ligand-induced osteoclast differentiation via a stromal cell independent mechanism involving c-Jun repression. Proc. Natl. Acad. Sci. U. S. A 97, 7829–7834 (2000).

30 Srivastava, S. et al. Estrogen decreases osteoclast formation by down-regulating receptor activator of NF-kappa B ligand (RANKL)-induced JNK activation. J. Biol. Chem 276, 8836–8840 (2001).

31 Robinson, L. J. et al. Estrogen inhibits RANKL-stimulated osteoclastic differentiation of human monocytes through estrogen and RANKL-regulated interaction of estrogen receptor-alpha with BCAR1 and Traf6. Exp Cell Res 315, 1287–1301 (2009). 10.1016/j.yexcr.2009.01.014

32 Nakamura, T. et al. Estrogen prevents bone loss via estrogen receptor alpha and induction of Fas ligand in osteoclasts. Cell 130, 811–823 (2007).

33 Krum, S. A. et al. Estrogen protects bone by inducing Fas ligand in osteoblasts to regulate osteoclast survival. EMBO J 27, 535–545 (2008).

34 Saintier, D. et al. Estradiol inhibits adhesion and promotes apoptosis in murine osteoclasts in vitro. J Steroid Biochem. Mol. Biol 99, 165–173 (2006).

35 Park, H. et al. Interaction of Fas ligand and Fas expressed on osteoclast precursors increases osteoclastogenesis. J Immunol 175, 7193–7201 (2005).

36 Kovacic, N. et al. The Fas/Fas ligand system inhibits differentiation of murine osteoblasts but has a limited role in osteoblast and osteoclast apoptosis. J Immunol 178, 3379–3389 (2007).

37 Kovacic, N. et al. Fas receptor is required for estrogen deficiency-induced bone loss in mice. Lab Invest 90, 402–413 (2010). 10.1038/labinvest.2009.144

38 Wang, L. et al. Osteoblast-induced osteoclast apoptosis by fas ligand/FAS pathway is required for maintenance of bone mass. Cell Death Differ 22, 1654–1664 (2015). 10.1038/cdd.2015.14

39 Warren, A. et al. The NAD salvage pathway in mesenchymal cells is indispensable for skeletal development in mice. Nat Commun 14, 3616 (2023). 10.1038/s41467-023-39392-7

40 Bertels, J. C., He, G. & Long, F. Metabolic reprogramming in skeletal cell differentiation. Bone Res 12, 57 (2024). 10.1038/s41413-024-00374-0

41 Kushwaha, P. et al. Mitochondrial fatty acid beta-oxidation is important for normal osteoclast formation in growing female mice. Front Physiol 13, 997358 (2022). 10.3389/fphys.2022.997358

42 Song, C. et al. Sexual dimorphism of osteoclast reliance on mitochondrial oxidation of energy substrates in the mouse. JCI Insight 8 (2023). 10.1172/jci.insight.174293

43 Li, B. et al. Both aerobic glycolysis and mitochondrial respiration are required for osteoclast differentiation. FASEB J 34, 11058–11067 (2020). 10.1096/fj.202000771R

44 Cambronne, X. A. et al. Biosensor reveals multiple sources for mitochondrial NAD(+). Science 352, 1474-1477 (2016). 10.1126/science.aad5168

45. Almeida, M. & Porter, R. M. Sirtuins and FoxOs in osteoporosis and osteoarthritis. Bone 121, 284-292 (2019). 10.1016/j.bone.2019.01.018

46 Wang, C. et al. Poly-ADP-ribosylation-mediated degradation of ARTD1 by the NLRP3 inflammasome is a prerequisite for osteoclast maturation. Cell Death Dis 7, e2153 (2016). 10.1038/cddis.2016.58

47 Moon, Y. J. et al. Sirtuin 6 in preosteoclasts suppresses age- and estrogen deficiency-related bone loss by stabilizing estrogen receptor alpha. Cell Death Differ 26, 2358–2370 (2019). 10.1038/s41418-019-0306-9

48 Gomes, A. P. et al. Declining NAD(+) Induces a Pseudohypoxic State Disrupting Nuclear-Mitochondrial Communication during Aging. Cell 155, 1624–1638 (2013). S0092-8674(13)01521-3 [pii];10.1016/j.cell.2013.11.037 [doi]

49 Zhang, H. et al. NAD(+) repletion improves mitochondrial and stem cell function and enhances life span in mice. Science 352, 1436–1443 (2016). 10.1126/science.aaf2693

50 Guarente, L., Sinclair, D. A. & Kroemer, G. Human trials exploring anti-aging medicines. Cell Metab 36, 354–376 (2024). 10.1016/j.cmet.2023.12.007

51 Rongvaux, A. et al. Nicotinamide phosphoribosyl transferase/pre-B cell colony-enhancing factor/visfatin is required for lymphocyte development and cellular resistance to genotoxic stress. J Immunol 181, 4685–4695 (2008). 10.4049/jimmunol.181.7.4685

52 Martin-Millan, M. et al. The estrogen receptor-alpha in osteoclasts mediates the protective effects of estrogens on cancellous but not cortical bone. Mol Endocrinol 24, 323–334 (2010). 10.1210/me.2009-0354

53 Bouxsein, M. L. et al. Guidelines for assessment of bone microstructure in rodents using micro-computed tomography. J Bone Miner Res 25, 1468–1486 (2010). 10.1002/jbmr.141

54 Dempster, D. W. et al. Standardized nomenclature, symbols, and units for bone histomorphometry: a 2012 update of the report of the ASBMR Histomorphometry Nomenclature Committee. J Bone Miner Res 28, 2–17 (2013). 10.1002/jbmr.1805

55 Law, C. W., Chen, Y., Shi, W. & Smyth, G. K. voom: Precision weights unlock linear model analysis tools for RNA-seq read counts. Genome Biol 15, R29 (2014). 10.1186/gb-2014-15-2-r29

56 Bartell, S. M. et al. FoxO proteins restrain osteoclastogenesis and bone resorption by attenuating H2O2 accumulation. Nat Commun 5, 3773 (2014). 10.1038/ncomms4773

